# Chemogenetic activation of microglial Gi signaling decreases microglial surveillance and impairs neuronal synchronization

**DOI:** 10.1101/2024.02.12.579861

**Authors:** Shunyi Zhao, Lingxiao Wang, Yue Liang, Jiaying Zheng, Anthony D. Umpierre, Long-Jun Wu

## Abstract

Microglia actively survey the brain and dynamically interact with neurons to maintain brain homeostasis. Microglial Gi-protein coupled receptors (Gi-GPCRs) play a critical role in microglia-neuron communications. However, the impact of temporally activating microglial Gi signaling on microglial dynamics and neuronal activity in the homeostatic brain remains largely unknown. In this study, we employed Gi-based Designer Receptors Exclusively Activated by Designer Drugs (Gi-DREADD) to selectively and temporally modulate microglial Gi signaling pathway. By integrating this chemogenetic approach with *in vivo* two-photon imaging, we observed that exogenous activation of microglial Gi signaling transiently inhibited microglial process dynamics, reduced neuronal activity, and impaired neuronal synchronization. These altered neuronal functions were associated with a decrease in interactions between microglia and neuron somata. Altogether, this study demonstrates that acute, exogenous activation of microglial Gi signaling can regulate neuronal circuit function, offering a potential pharmacological target for neuromodulation through microglia.

## Introduction

Microglia actively survey the brain through their dynamic processes.^1–3^ Their surveillance and interactions with neurons have been linked to neuronal activity changes.^4, 5^ Specifically, physical microglia-neuron interactions have been associated with dampening hyperactive neurons, promoting hypoactive neuronal activity, or maintaining the hypoactive state of neurons.^6–9^ These functions are partially mediated by microglial Gi-protein coupled receptors (Gi-GPCRs), such as the P2Y12 receptor.^6, 8, 9^ Genetic knockout of the P2Y12 receptor in microglia has been shown to elevate anxiety, disrupt general anesthesia, and increase seizure severity in mice.^6, 8–12^ Additionally, microglial-specific Gi signaling inhibition through pertussis toxin expression (PTX) decreased microglial surveillance and increased neuronal activity.^13^ These findings indicate that microglial Gi-GPCRs are crucial in regulating neuronal activity and emerge as intriguing targets for neuromodulation. However, the impact of specifically activating microglial Gi signaling on the neural circuit function remains unknown.

To selectively activate microglial Gi signaling, recent studies employed a chemogenetic tool, Designer Receptors Exclusively Activated by Designer Drugs (DREADD).^14^ Utilizing these engineered Gi-GPCRs, we demonstrated that microglial Gi-DREADD activation could mimic endogenous Gi-mediated chemotaxis.^15^ Furthermore, this method has been successfully applied in disease conditions, such as pain and epilepsy, to explore function of microglial Gi signaling.^16^ Acute activation of microglial Gi signaling after seizure onset increased microglia-neuron soma interactions, altered microglial morphology, and decreased seizure severity.^17^ Chronic activation of microglial Gi signaling after peripheral nerve injury decreased the inflammatory cytokine release and ameliorated chronic pain.^18, 19^ These results indicate the potential of microglial Gi signaling to relieve pathological neuron hyperactivity.

In the current study, we used two-photon imaging and microglial chemogenetics to investigate how microglial Gi signaling influences microglia-neuron interaction and neural circuit function in the homeostatic brain. We observed that microglia retracted their processes after Gi-DREADD activation. This morphology alteration led to decreased interactions with neuronal soma, which was associated with reduced neuronal activity and impaired synchronization. Our findings address an intriguing question of how microglia are integrated into neuronal circuits. Regulation of neural circuit function through microglial Gi signaling could provide a new pharmacological target for neuromodulation.

## Results

### Chemogenetic activation of microglial Gi signaling decreases microglial process surveillance

Genetically inhibiting microglial Gi signaling by PTX decreases microglial surveillance.^13^ To activate Gi signaling in microglia, we induced hM4Di (Gi-DREADD) expression in microglia using a *Tmem119^CreER/+^:R26^hM4Di/+^:Cx3cr1^GFP/+^*mouse line (Fig. 1A). After tamoxifen treatment, antibody detection of the hemagglutinin (HA) tag fused to the Gi-DREADD receptor was specifically localized to the membrane of GFP^+^ microglia, but not in CD206^+^ border associated macrophages (BAM), NeuN^+^ neurons, GFAP^+^ astrocytes, or CC1^+^ oligodendrocytes (Fig. 1B and C). We then administered the DREADD agonist Deschloroclozapine (DCZ, *i.p.*) to activate microglial Gi-DREADD.^20^ A low dose of DCZ (100 μg/kg) takes effect in the brain within 30 minutes and is eliminated from the body within 24 hours.^20^

**Figure 1.**
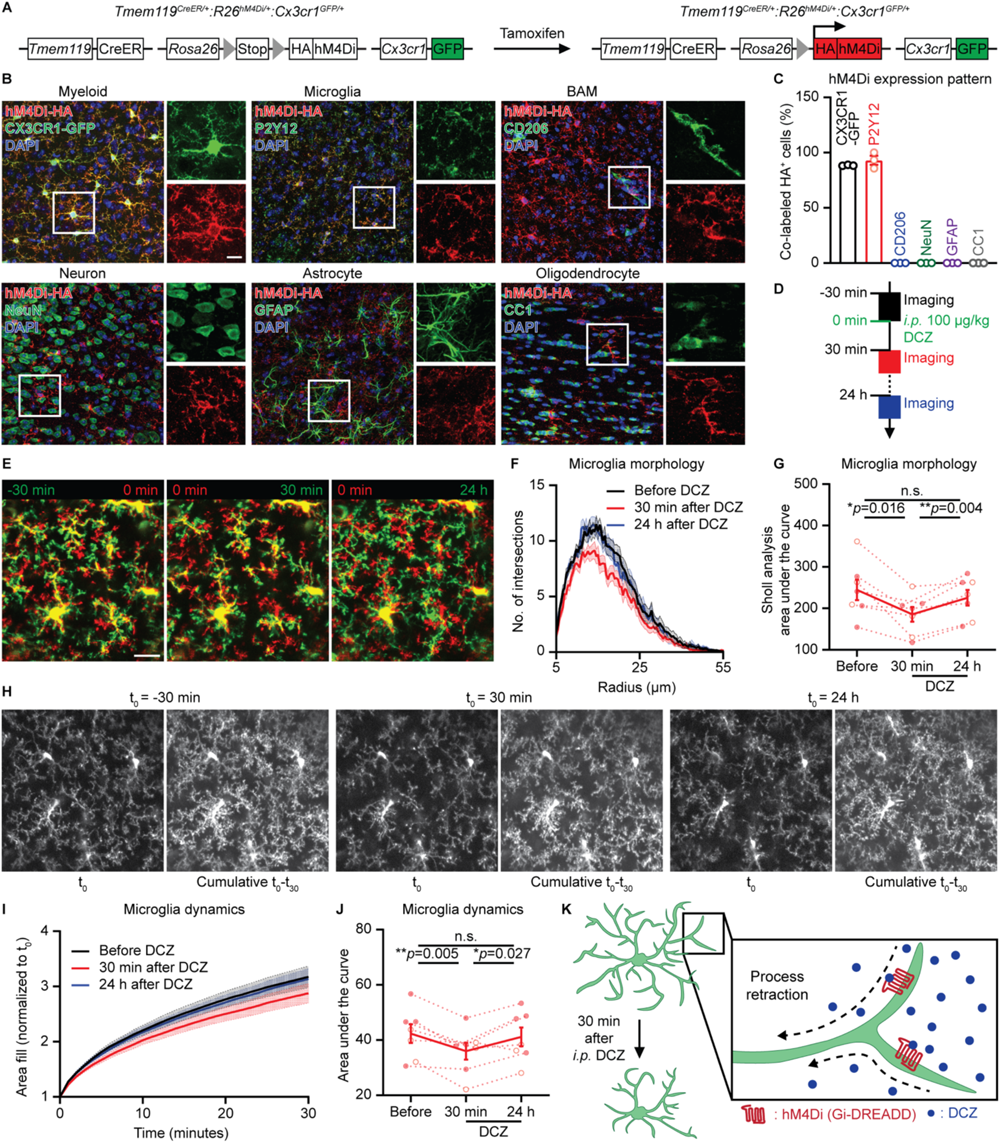
Microglial Gi signaling reduces microglial process surveillance. (**A**) The genetic strategy employed in *Tmem119^CreER/+^:R26^hM4Di/+^:Cx3cr1^GFP/+^* mice for the expression of hM4Di (Gi-DREADD) and GFP in microglia. (**B** and **C**) Immunostaining (B) and quantification (C) reveal the membrane localization of HA-tagged hM4Di (red) exclusively on microglia (GFP^+^ or P2Y12^+^, green), not on border-associated macrophages (BAM; CD206^+^, green), neurons (NeuN^+^, green), astrocytes (GFAP^+^, green), and oligodendrocytes (CC1^+^, green). Therefore, Gi-DREADD is predominantly expressed by microglia in the brain of *Tmem119^CreER/+^:R26^hM4Di/+^:Cx3cr1^GFP/+^* mice (N=3 mice in each group). Scale bar: 10 μm (B). (**D**) Timeline of *in vivo* two-photon imaging. 30-minute videos were recorded before, 30 minutes after, and 24 hours after 100 μg/kg DCZ *i.p.* administration. (**E**) Representative two-photon images illustrate microglial morphology alterations in *Tmem119^CreER/+^:R26^hM4Di/+^:Cx3cr1^GFP/+^*mice at each study phase. Increased red signal area in the mid-panel indicates microglial process retraction 30 minutes following DCZ injection. Left: before (green: -30 min; red: 0 min). Mid: 30 minutes after (red: 0 min; green: 30 min). Right: 24 h after (red: 0 min; green: 24 h). Scale bar: 20 μm. (**F** and **G**) Sholl analysis (F) and the area under the curve quantification (G) demonstrate microglial process retraction 30 minutes after DCZ administration, with morphology returning to baseline 24 hours later in *Tmem119^CreER/+^:R26^hM4Di/+^:Cx3cr1^GFP/+^*mice (30 microglia from N=7 mice in each group). (**H**) Two-photon images show cumulative microglial surveillance before, 30 minutes after, and 24 hours after DCZ administration in *Tmem119^CreER/+^:R26^hM4Di/+^:Cx3cr1^GFP/+^*mice. Microglia exhibited reduced surveillance area post-injection. Upper: microglial surveillance area at the beginning of each study phase. Lower: 30-minute cumulative microglial surveillance area of each study phase. Scale bar: 20 μm. (**I** and **J**) Normalized cumulative microglial surveillance area (I) and the area under the curve quantification (J) demonstrate decreased microglial surveillance 30 minutes after DCZ administration, returning to baseline 24 hours later in *Tmem119^CreER/+^:R26^hM4Di/+^:Cx3cr1^GFP/+^*mice (N=7 mice in each group). (**K**) Diagram illustrating microglial Gi-DREADD activation leads to microglial process retraction and decreased surveillance 30 minutes after DCZ administration. In all graphs, each point indicates an individual mouse (solid dots: male mice; hollow dots: female mice). Data are represented as the mean ± SEM. Repeated measures one-way ANOVA followed by a Bonferroni’s *post hoc* test (G and J). n.s.: not significant. \**p*<0.05 and \*\**p*<0.01. The exact *p* values are directly provided in the figure panels.

We used *in vivo* two-photon microscopy to image microglial dynamics in the cortex following DCZ Gi-DREADD activation in awake mice. Surprisingly, we found that systemic DCZ acutely reduced microglia process dynamics (Fig. 1D-G, and Video S1). DCZ-induced microglial process retraction was gradually returned to baseline levels 24 hours later (Fig. 1D-G). Furthermore, analyses of process dynamics over 30 minutes showed that microglia survey less area of the cortical parenchyma 30-60 minutes after DCZ injection, compared with baseline and 24 hours after DCZ injection (Fig. 1H-J). Together, these results indicate that chemogenetic activation of microglial Gi signaling transiently decreases microglial process surveillance in awake mice (Fig. 1K). As a control, in *Tmem119^CreER/+^:Cx3cr1^GFP/+^* mice (Fig. S1A), DCZ injection did not alter microglia morphology (Fig. S1B-D) and surveillance area (Fig. S1E-G). Similarly, saline injection in *Tmem119^CreER/+^:R26^hM4Di/+^:Cx3cr1^GFP/+^*mice (Fig. S1H and I) also failed to change microglia morphology (Fig. S1J-L) and surveillance area (Fig. S1M-O).

Neuronal activity alters microglial surveillance.^6, 12, 21–23^ It is possible that DCZ injection indirectly modulates microglial morphology by affecting neuronal activity. To test this possibility, we imaged microglia dynamics in response to DCZ in *Tmem119^CreER/+^:R26^hM4Di/+^:Cx3cr1^GFP/+^*mice under anesthesia (Fig. S2A). Given that microglia increase surveillance during the first 20 minutes of anesthesia,^22, 23^ we initiated imaging microglia 20 minutes after the anesthesia induction followed by DCZ injection (Fig. S2B). Similar to observations in the awake state, chemogenetic activation of microglial Gi signaling induced microglia process retraction (Fig. S2C-E, and Video S2) and decreased microglial process surveillance in anesthetized mice (Fig. S2F-H). In control mice lacking Gi-DREADD expression (Fig. S2I), DCZ injection did not affect microglia morphology (Fig. S2J-L) or surveillance area (Fig. S2M-O). These results support the direct role of microglia Gi signaling in reducing microglia process dynamics.

Two-photon microscopy often limits imaging depth to the cortex. We additionally wanted to assess more universal changes in microglial morphology across brain regions following Gi-DREADD activation. To evaluate microglial morphology, we performed Iba1 immunostaining and then utilized trainable Weka Segmentation to render microglia for Sholl analyses (Fig. 2A).^24^ Our results suggest microglia decrease their process length and exhibited less complexity in the cortex (Fig. 2B-D), corpus callosum (Fig. 2E-G), and hippocampus (Fig. 2H-J) when DCZ is administered to Gi-DREADD mice (versus their *Tmem119^CreER/+^* genotype controls). In contrast, saline injection did not alter microglial morphology (Fig. S3A-I). Thus, activation of microglial Gi signaling induces process retraction across brain regions.

**Figure 2.**
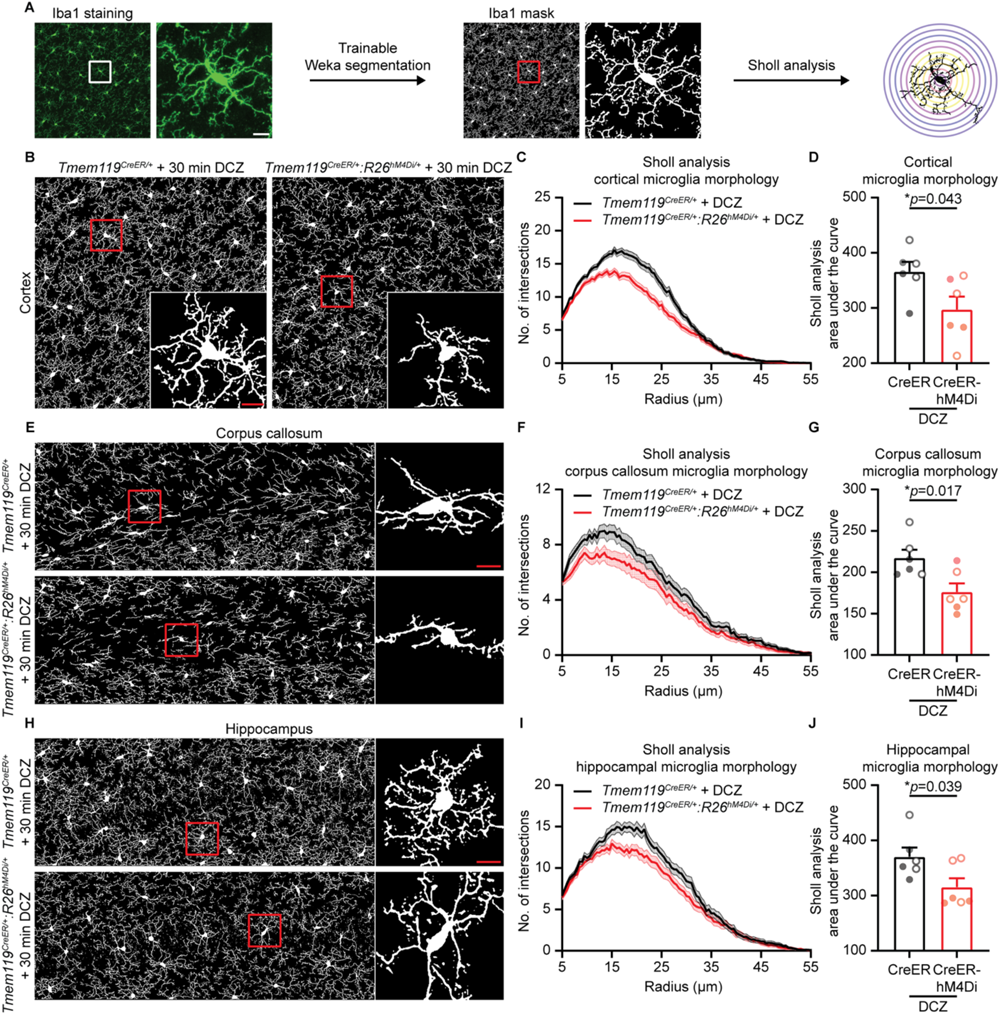
Activating microglial Gi-DREADD reduces microglia process length in various brain regions. **(A)** Diagram illustrating the data processing. Iba1 immunostaining images were thresholded using trainable Weka segmentation, and microglia were manually selected for Sholl analysis. (**B**-**D**) Analysis of cortical microglia in DCZ-administered control and *Tmem119^CreER/+^:R26^hM4Di/+^* mice. Representative Iba1 mask (B), Sholl analysis (C), and the area under the curve quantification (D) demonstrate process retraction in cortical microglia in *Tmem119^CreER/+^:R26^hM4Di/+^* mice (90 microglia from N=6 mice in each group). (**E**-**G**) Analysis of microglia in the corpus callosum. Representative Iba1 mask (E), Sholl analysis (F), and the area under the curve quantification (G) reveal process retraction in corpus callosum microglia of DCZ-administered *Tmem119^CreER/+^:R26^hM4Di/+^*mice (60 microglia from N=6 mice in each group). (**H**-**J**) Evaluation of hippocampal microglia. Representative Iba1 mask (H), Sholl analysis (I), and the area under the curve quantification (J) show process retraction in hippocampal microglia 30 minutes after DCZ administration in *Tmem119^CreER/+^:R26^hM4Di/+^* mice (60 microglia from N=6 mice in each group). In all graphs, each point indicates an individual mouse (solid dots: male mice; hollow dots: female mice). Data are represented as the mean ± SEM. Unpaired *t*-test (D, G, and J). \**p*<0.05. The exact *p* values are directly provided in the figure panels. Scale bar: 10 μm (A, B, E, and H).

### Activation of microglial Gi signaling reduces microglia-neuron soma interactions

Microglia engage in dynamic intercellular communication through close contact and surveillance of neuronal somata.^7, 25^ Therefore, we investigated the impact of Gi-DREADD-mediated microglial process retraction on microglia-neuron soma interactions. To visualize these interactions in real-time, we injected AAV2/5-CaMKIIa-tdTomato to label excitatory neurons in both *Tmem119^CreER/+^:Cx3cr1^GFP/+^*and *Tmem119^CreER/+^:R26^hM4Di/+^:Cx3cr1^GFP/+^*mice (Fig. 3A). We then acquired videos before, 0-60 minutes after, and 24 hours after the DCZ injection (Fig. 3B). The DCZ injection in *Tmem119^CreER/+^:Cx3cr1^GFP/+^*control mice had no significant effect on microglia-neuron soma interactions (Fig. 3C). However, in *Tmem119^CreER/+^:R26^hM4Di/+^:Cx3cr1^GFP/+^*mice, a rapid reduction in microglia surveillance specifically decreases process contact frequency with neuronal somata 30-60 minutes after DCZ administration (Fig. 3D, E, and Video S3). However, for processes remaining in contact with somata, their duration of interaction was similar to baseline (Fig. 3F). This suggests that certain stable physical interactions might be formed, which appear to be resistant to microglial Gi-DREADD activation. Additionally, somatic interaction events by microglia returned to the baseline levels 24 hours post-DCZ injection (Fig. 3E).

**Figure 3.**
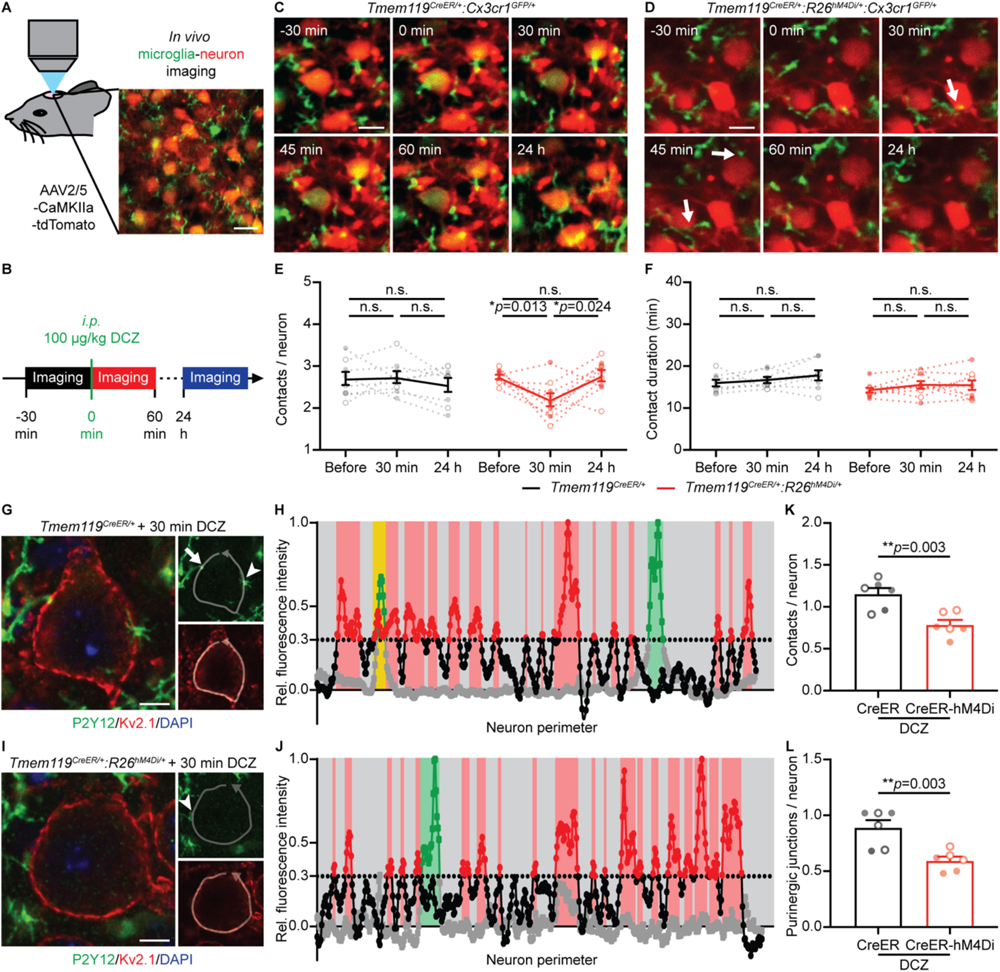
Chemogenetic activation of microglial Gi signaling decreases microglia-neuron soma interactions. **(A)** Experimental design diagram illustrating two-photon imaging of GFP^+^ microglia (green) and tdTomato^+^ excitatory neurons (red) for visualizing microglia-neuron soma interactions in real-time. Scale bar: 20 μm. **(B)** Timeline of *in vivo* two-photon imaging to assess the microglia-neuron soma interactions. Videos were recorded before, 0-60 minutes after, and 24 hours after 100 μg/kg *i.p.* DCZ administration. (**C** and **D**) Time-lapse images of microglia (green) and neuron soma (red) interactions at each study phase in *Tmem119^CreER/+^:Cx3cr1^GFP/+^*(C) and *Tmem119^CreER/+^:R26^hM4Di/+^:Cx3cr1^GFP/+^*mice (D). Arrows indicate the retracting processes from neuronal soma (D). Scale bar: 10 μm. (**E** and **F**) Quantifications of microglia contact number (E) and contact duration (F) reveal fewer contact events 30-60 minutes after DCZ administration in *Tmem119^CreER/+^:R26^hM4Di/+^:Cx3cr1^GFP/+^* mice, while the contact duration was unchanged. The microglia-neuron soma interaction number returned to baseline 24 hours following DCZ injection (103 neurons from N=8 mice, *Tmem119^CreER/+^:Cx3cr1^GFP/+^*;133 neurons from N=9 mice, *Tmem119^CreER/+^:R26^hM4Di/+^:Cx3cr1^GFP/+^*). (**G**-**J**) Immunostaining (G and I) and plots of neuron perimeter versus fluorescence intensity (H and J) displaying neuronal Kv2.1 membrane domains (Kv2.1^+^ and P2Y12^neg^, red), non-purinergic microglial contacts (Kv2.1^neg^ and P2Y12^+^, green), and purinergic junctions (Kv2.1^+^ and P2Y12^+^, yellow). 30 minutes after DCZ injection, neurons had fewer microglial contacts and purinergic junctions in *Tmem119^CreER/+^:R26^hM4Di/+^* mice. The arrow indicates the purinergic junction (G), and the arrowheads indicate the non-purinergic contacts (G and I). Scale bar: 5 μm. (**K** and **L**) Quantifications reveal microglia processes had fewer total contacts (K) and purinergic junctions (L) per neuron 30 minutes after DCZ administration in *Tmem119^CreER/+^:R26^hM4Di/+^* mice (300 neurons from N=6 mice in each group). In all graphs, each point indicates an individual mouse (solid dots: male mice; hollow dots: female mice). Data are represented as the mean ± SEM. Repeated measures one-way ANOVA followed by a Bonferroni’s *post hoc* test (E and F). Unpaired *t*-test (K and L). n.s.: not significant. \**p*<0.05 and \*\**p*<0.01. The exact *p* values are directly provided in the figure panels.

Somatic contacts between microglia processes and neuronal somata have been previously characterized as purinergic junctions, involving microglial P2Y12 receptors and neuronal Kv2.1^+^ membrane domains.^7, 26^ We performed immunostaining of P2Y12 and Kv2.1 to assess the interaction 30 minutes after the DCZ injection (Fig. 3G and I). Relative fluorescence intensity around neuronal perimeter quantified neuronal Kv2.1 membrane domains (Kv2.1≥0.3 and P2Y12:<0.3, red), non-purinergic microglial contacts (Kv2.1:<0.3 and P2Y12≥0.3, green), and purinergic junctions (Kv2.1≥0.3 and P2Y12≥0.3, yellow) (Fig. 3H and J). We found that both the total microglia-neuron soma interaction and the formation of purinergic junctions decreased in *Tmem119^CreER/+^:R26^hM4Di/+^*mice after DCZ treatment (Fig. 3K and L). No differences were observed in microglial-neuron soma interaction in the saline-injected *Tmem119^CreER/+^* and *Tmem119^CreER/+^:R26^hM4Di/+^*mice (Fig. S3J-O). These results suggest the microglial Gi signaling-induced process retraction further disrupts microglial interactions with neuronal somata.

### Activation of microglial Gi signaling reduces neuronal activity and synchronization

Physical microglia-neuron interactions are closely linked to the regulation of neuronal activity.^2–5^ Recent findings have indicated that purinergic junctions formed between microglia processes and neuron somata serve a protective role against neuronal excitotoxicity in stroke.^26^ Here, we employed *in vivo* two-photon calcium imaging (AAV9-CaMKIIa-GCaMP6s) to assess neuronal activity following microglial Gi-DREADD activation (Fig. 4A and B). In *Tmem119^CreER/+^* control mice, DCZ did not alter neuronal activity (Fig. 4C). However, in *Tmem119^CreER/+^:R26^hM4Di/+^*mice, we observed a reduction in neuronal active time, signal area, and calcium transient events 30 minutes after the DCZ injection (Fig. 4D-G), while the duration of calcium transients remained unaltered (Fig. 4H). Next, we investigated the local neuronal network synchronization using the MATLAB-based Mic2net program.^27^ Interestingly, we found a decrease in neuronal connectivity and reduced synchronization 30 minutes after DCZ injection (Fig. 4I-L). Reduced neuronal activity and synchronization returned to baseline levels 24 hours after DCZ administration (Fig. 4E-G, K, and L). These data indicate that temporary activation of microglial Gi signaling reduces neuronal activity and synchronization.

**Figure 4.**
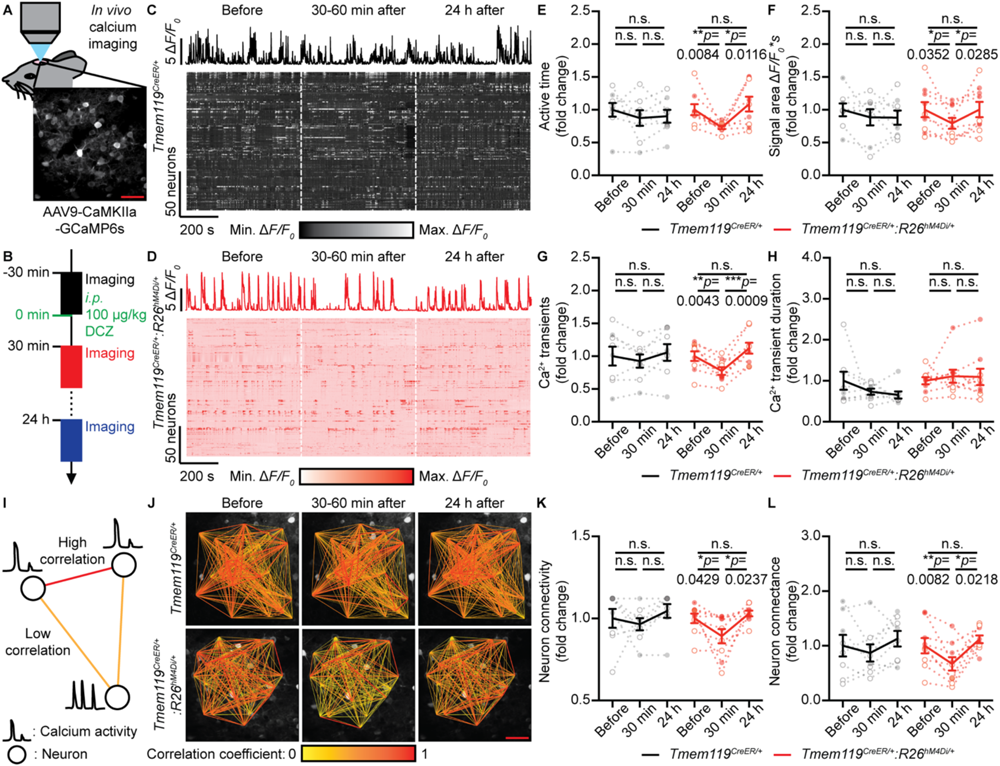
Microglial Gi-DREADD activation reduces neuron activity and synchronization. **(A)** Experimental design of two-photon imaging for assessing excitatory neuron activity. Scale bar: 50 μm. **(B)** Timeline of *in vivo* two-photon imaging to evaluate the neuronal activity. Videos were recorded before, 30-60 minutes after, and 24 hours after 100 μg/kg *i.p.* DCZ administration. (**C** and **D**) Single neuronal calcium activity traces (upper) and heatmap of all neuronal calcium activity (lower) before, 30-60 minutes after, and 24 hours after DCZ administration in *Tmem119^CreER/+^* (C) and *Tmem119^CreER/+^:R26^hM4Di/+^* mice (D). Reduced neuronal activity was observed 30-60 minutes after DCZ injection in *Tmem119^CreER/+^:R26^hM4Di/+^*mice, recovering 24 hours later. (**E**-**H**) Quantifications revealing decreased neuronal active time (E), signal area (F), and calcium transient events (G) 30 minutes after the DCZ injection in *Tmem119^CreER/+^:R26^hM4Di/+^* mice. The downregulations recovered to baseline 24 hours post-injection. The calcium transient duration was not affected by DCZ administration (H, 155 neurons from N=8 mice, *Tmem119^CreER/+^*; 190 neurons from N=9 mice, *_Tmem119_CreER/+_:R26_hM4Di/+*_)._ (**I**) Diagram showing the analysis of intercellular calcium activity correlation coefficient (high, red; low, yellow) to determine neuronal network synchronization. (**J**) Representative neuronal network analysis plotted on top of two-photon calcium images at each study phase in *Tmem119^CreER/+^* and *Tmem119^CreER/+^:R26^hM4Di/+^* mice. Fewer red edges in the lower mid-panel suggest the neuronal hyposynchronization 30 minutes after DCZ injection in *Tmem119^CreER/+^:R26^hM4Di/+^* mice. Scale bar: 50 μm. (**K** and **L**) Quantifications revealing decreased neuron connectivity (K) and neuron connectance (L) 30 minutes after the DCZ injection. The reductions returned to baseline 24 hours after the DCZ injection in *Tmem119^CreER/+^:R26^hM4Di/+^* mice (N=8 mice, *Tmem119^CreER/+^*; N=9 mice, *Tmem119^CreER/+^:R26^hM4Di/+^*). In all graphs, each point indicates an individual mouse (solid dots: male mice; hollow dots: female mice). Data are represented as the mean ± SEM. Repeated measures one-way ANOVA followed by a Bonferroni’s *post hoc* test (E-H, K, and L). n.s.: not significant. \**p*<0.05, \*\**p*<0.01 and \*\*\**p*<0.001. The exact *p* values are directly provided in the figure panels.

Despite Tmem119 being predominantly expressed in microglia in the central nervous system, its expression in peripheral cell types necessitated additional verification that the observed reductions in neuronal activity and synchronization were indeed caused by microglial Gi signaling (Fig. S4A and B). The query of the RNA sequencing database demonstrates that the combination of genetic microglia manipulation (Tmem119 driven) and pharmacological microglia ablation (targeting CSF1R) effectively narrowed the cell population to microglia (Fig. S4B). Therein, we ablated microglia through PLX3397 administration, a CSF1R inhibitor, to alleviate the off-target concern. If the reduced neuronal activity after DCZ injection was dependent on microglia, we should rescue the phenotype after microglia ablation. Indeed, when PLX3397 chow was provided 2 weeks before the calcium imaging (Fig. S4C-E), we did not observe the previously noted reductions in neuronal active time, signal area, and calcium transient events (Fig. S4F-J). Additionally, there were no alterations in the local neuronal connectivity and synchronization 30 minutes following the DCZ injection (Fig. S4K-M). These results strongly support that activation of microglia Gi signaling truly reduced neuron activity and synchronization.

## Discussion

In this study, we combined microglial chemogenetics with *in vivo* two-photon imaging to investigate the impact of microglial Gi signaling in the regulation of neuronal function. Upon activation of microglial Gi signaling, we observed decreased microglial surveillance, reduced neuronal activity, and impaired local neuronal synchrony. The altered neuronal functions were associated with a decrease in microglial process interaction with neuronal somata. Collectively, these findings suggest that microglia Gi signaling can transiently modulate neuronal activity through physical interactions with neurons in the homeostatic brain.

### Regulation of microglial process dynamics by Gi signaling pathway

Microglia express a number of Gi-GPCRs, such as CX3CR1, P2Y12, and C3aR.^28^ Previous studies have shown that the microglia Gi signaling pathway governs the microglial process functions, including chemotaxis and dynamic movement. Specifically, P2Y12 and C3aR are known to mediate microglial process chemotaxis.^29, 30^ The absence of CX3CR1 leads to reduced microglia process length.^31^ Furthermore, suppressing Gi signaling in microglia via selective PTX expression resulted in shorter and less dynamic microglial processes.^13^ These varied consequences of inhibiting microglial Gi-GPCR underscore the importance of the Gi signaling pathway in regulating microglial process morphology and motility.

In our study, we utilized chemogenetic manipulation to selectively activate microglial Gi signaling, which leads to a reduction in microglial process length and a decrease in surveillance. Additionally, this effect is transient, with microglial morphology and surveillance returning to baseline levels 24 hours after DCZ injection. Together with previous reports, these observations suggest a “U-shaped” regulatory function by the microglial Gi signaling on microglia dynamics, where both increased and decreased Gi signaling can lead to lower microglial surveillance.

The microglial morphology alteration may stem from the reorganization of the cytoskeleton, involving filamentous actin (F-actin) or microtubules. Activation of Gi-GPCR results in cyclic AMP (cAMP) downregulation, which leads to the decompartmentalization of F-actin and the subsequent collapse of microglial filopodia.^32^ Additionally, Gi-GPCR is implicated in the microtubule dynamics.^33^ Small Gi proteins are able to inhibit microtubule assembly by interacting with the GTP cap of growing microtubules,^34^ offering an alternative mechanism by which Gi-DREADD activation could attenuate microglia surveillance. However, these hypotheses need further experiments to confirm.

Last but not least, as of now, there are no established methods for selectively and temporally manipulating microglia morphology *in vivo*. Our findings that microglial Gi-DREADD activation can transiently decrease surveillance may open up a new avenue for understanding the function of microglial dynamics in neural circuits.

### Physical microglia-neuron interactions in the regulation of neuronal functions

Previous studies have demonstrated that genetic knockout of endogenous microglial Gi-GPCRs, like P2Y12, leads to increased anxiety and decreased seizure severity in mice.^6, 10–12, 35^ Moreover, inhibition of Gi signaling in microglia has been associated with increased neuronal activity and synchronization.^13^ In line with these findings, our current study shows that the chemogenetic activation of microglial Gi signaling reduces neuronal activity and disrupts the local synchronization of neural networks. This is supported by a recent study showing microglial Gi-DREADD activation induces neuronal hypoactivity, decreases norepinephrine release and promotes sleep.^36^ Thus, microglial Gi signaling is able to suppress neuronal activity in the homeostatic brain.

Reduced neuronal activity after microglial Gi activation was correlated with fewer interactions between microglia processes and neuronal somata. These interactions are thought to represent a novel method of neuromodulation.^4, 5^ For instance, during a stroke or seizure, microglial processes could be attracted by the overactive neuron, forming somatic interactions to prevent excitotoxicity.^12, 26^ Interestingly, our findings imply that in a non-pathological state, microglia-neuron interactions might actually enhance neuronal activity. Consistent with this, our recent study demonstrated that microglia extended their processes to shield inhibitory synapses on neuronal somata, thereby promoting post-anesthesia neuronal activity.^7^ Therefore, it is possible that microglial Gi-DREADD-mediated surveillance downregulation could disrupt the shielding of inhibitory synapses on neuronal somas, leading to reduced neuronal activity.

Beyond these microglia-neuron soma interactions, microglia are also known to contact with excitatory synapses. An earlier study has shown that microglial contacts with dendritic spines increase spine calcium activity, thereby enhancing the activity of the local neuronal network.^37^ Thus, reduced microglial surveillance might also decrease interactions with excitatory synapses, potentially reducing neuronal activity.

### Microglial Gi signaling as a potential target for neuromodulation in health and disease

Recent studies have highlighted the integral role of microglia in the regulation of neuronal activity and synaptic plasticity, such as learning and memory,^38, 39^ sleep,^40–42^ anxiety-like behaviors,^10^ compulsive behaviors,^43^ alcohol intake,^44^ pain,^45^ and seizures.^46^ Beyond direct physical interactions with neurons, these studies suggest alternative regulatory mechanisms whereby microglia release soluble factors and phagocytose extracellular matrix (ECM) to modulate neuronal activity.^47^ Microglia Gi-GPCRs were reported to participate in the regulation of neuronal functions. For example, pharmacologically inhibiting P2Y12 has been shown to prevent ketamine-induced ECM loss.^48^ Thus, the activation of microglial Gi-DREADD might share similar mechanisms to modulate neuronal activity.

While Gi signaling canonically suppresses intracellular cAMP levels and inhibits associated pathways, emerging evidence indicates that microglial Gi signaling can also elevate microglial calcium activity.^36, 49^ Interestingly, microglial calcium activity is attuned to neuronal activity,^50^ and is implicated in the release of soluble factors^51, 52^ and in enhancing phagocytosis.^53, 54^ Therefore, Gi signaling-mediated calcium elevation might trigger the release of soluble factors and facilitate the phagocytosis of ECM to modulate neuronal activity and network synchronization.

Moreover, microglia Gi signaling can also be potential targets for neuromodulation in neurological diseases. Recent findings indicate that chemogenetic activation of microglia Gi-GPCRs could ameliorate seizures and relieve pain.^16, 55^ Chronic activation of Gi-DREADD has been shown to reduce the release of pro-inflammatory cytokines and alleviate neuropathic pain.^18, 19, 56, 57^ In the kainic acid (KA) induced seizure model, acute activation of microglial Gi-DREADD reduced neuronal hyperactivity, while chronic activation resulted in neuronal death, suggesting the balance maintained by microglia Gi signaling in the neuromodulation.^17^ Although further research is needed to elucidate the detailed mechanism by which microglial Gi signaling modulates neuronal function, our study identifies microglia Gi signaling as an intriguing target to fine-tune neuronal activity under physiological and pathological conditions.

## Methods

### Animals

The *Tmem119^CreER/CreER^* (C57BL/6-Tmem119^em^^1(cre/ERT^^2)Gfng^/J, 031820), *_R26_hM4Di/hM4Di* _(B6.129-*Gt(ROSA)26Sor*_*tm1(CAG-CHRM4*,-mCitrine)Ute*_/J, 026219), and_ *Cx3cr1^GFP/GFP^*(B6.129P2(Cg)-*Cx3cr1^tm1Litt^/*J, 005582) mouse lines were acquired from the Jackson Laboratory, and subsequently bred at Mayo Clinic. Tamoxifen (Sigma, 150 mg/kg, *i.p.*) was administered to all adult mice 4 times every 2 days to induce Cre-Lox recombination or serve as controls. All experiments were performed at least 4 weeks after tamoxifen injections. Both male and female mice were included in the studies. Mice were housed in a 12-hour light/dark cycle and climate-controlled environment, with *ad libitum* access to food and water. All experimental procedures were approved by the Mayo Clinic’s Institutional Animal Care and Use Committee (IACUC).

### Cranial window surgeries and virus injections

In preparation for cranial window surgeries, mice were provided with 0.2 mg/mL ibuprofen in their drinking water for 3 days prior. Anesthesia was induced with 3% isoflurane in oxygen, followed by maintenance at 1.5% in oxygen during surgery. Throughout the surgery, mice were kept on a heating pad. After cutting the skin to expose the skull, a craniotomy was performed to create a circular 4-mm-diameter window above the somatosensory cortex (centered at AP: -2.0; ML: +1.5). Subsequently, a sterile 5-mm glass coverslip (Warner Instruments) was implanted and affixed with dental cement (Tetric EvoFlow). The remaining skull was covered with iBond Total Etch glue (Heraeus) and cured with an LED Curing Light. Additional dental cement (Tetric EvoFlow) was again applied around the glass coverslip and cured. A custom-made head plate was then secured to the window using dental cement. Following the surgery, mice recovered under a heating lamp before being returned to their home cage, receiving 0.2 mg/mL ibuprofen in their drinking water for an additional 3 days.

For experiments requiring virus injections, after the craniotomy, AAVs were injected into the somatosensory cortex (AP: -2.0; ML: +1.5) using a microsyringe (Hamilton) and a glass pipette. During the virus injection, the exposed brain surface was kept moist with sterile phosphate-buffered saline (PBS). AAV2/5.CaMKII.tdTomato (Neurophotonics, 661-aav5) was used to label excitatory neurons and AAV9.CaMKII.GCaMP6s.WPRE.SV40 (Addgene, 107790) was employed to image excitatory neuronal calcium activity. 300 nL of AAVs were injected to target layer II/II neurons (DV: -0.3, 200 nL, 40 nL/min; DV: -0.2, 100 nL, 20 nL/min) with a 5-minute period for virus diffusion.

### Two-photon imaging

Prior to data acquisition, mice recovered for at least 4 weeks post-surgery and underwent a 3-day training period. During the training session, each mouse spent one hour per day on an air-lifted platform (NeuroTar) while being head-fixed under a two-photon microscope (Scientifica). The two-photon microscope was equipped with a Mai-Tai DeepSee laser (Spectra Physics) tuned to 940 nm. A 16x water immersion lens (Nikon) was utilized, and digital magnification (zoom 5-8) was applied based on experimental requirements. GFP and GCaMP6s signals passed through a 525/50 filter (Chroma), and tdTomato signal passed through a 630/75 filter (Chroma). Laser power was maintained at 4 mW or below.

Imaging in the somatosensory cortex was conducted 50-150 μm beneath the pial surface for microglia dynamics (20 μm thick z-stacked images, once per minute, 2 μm step size, 1024 × 1024 pixels) or 150-300 μm for imaging microglia-neuron interactions (20 μm Z-stacked images, 2 μm step size, once per minute, 1024 × 1024 pixels) and neuronal activity (1 Hz frame rate, 512 × 512 pixel resolution). Unless specifically stated, 30-minute videos were acquired before, 30-60 minutes, and 24 hours after 100 μg/kg DCZ injection (MedChemExpress). For imaging microglia dynamics in anesthetized mice, the mouse was induced with 3% isoflurane and was then maintained at 1.5% isoflurane for the anesthesia imaging. Under anesthesia, mice were placed on a heating pad (Physitemp) to maintain body temperature at 37°C. 20 minutes after the anesthesia induction, microglia dynamics images were acquired before and after DCZ injections.

### Immunofluorescence staining

Isoflurane anesthetized mice underwent transcardial perfusion with 40 mL PBS followed by 40 mL of cold 10% formaldehyde in PBS (LabChem). The whole brain was harvested, immersed in 10% formaldehyde overnight at 4 °C, and cryoprotected in 30% sucrose in PBS for a minimum 48 hours before cryosectioning. Free-floating sections (30 µm) were obtained using a cryostat (Leica). For immunostaining, sections were rinsed 3 times in the tris-buffered saline (TBS), blocked with TBS buffer containing 5% donkey serum and 0.4% Triton X-100 at room temperature for 1 hour. Subsequently, they were incubated overnight at 4 °C with primary antibodies: rabbit anti-HA (1:200, Cell Signaling Technology, 3724S), chicken anti-GFP (1:1000, Aves Labs, GFP-1010), rat anti-P2Y12 (1:100, BioLegnd, 848002), rat anti-CD206 (1:250, Bio-Rad, MCA2235), mouse anti-NeuN (1:500, Abcam, Ab104224), mouse anti-GFAP (1:500, Cell Signaling Technology, 3670S), mouse anti-CC1 (1:200, MilliporeSigma, OP80), rabbit anti-Iba1 (1:1000, Abcam, Ab178847), and mouse anti-Kv2.1 (1:500, antibodies-online Inc., ABIN1304762). Afterward, sections were washed with TBS 3 times and incubated with donkey secondary antibodies (1:500, Alexa-Fluor 488/594/647 anti-rabbit, anti-mouse, or anti-rat, Invitrogen, A21202, A21207, A21209, A31573, and A32787 or 1:500, Alexa-Fluor 488 anti-chicken, Jackson ImmunoResearch, 703-545-155) in blocking buffer for 120 min at room temperature. After 3 additional washes, sections were mounted with DAPI Fluoromount-G mounting medium (SouthernBiotech). Fluorescent images were acquired using a laser scanning confocal microscope (LSM 980, Zeiss). 10 μm Z-stacked images were obtained by a 20x, 40x, or 63x objective based on experimental requirements. Image brightness and contrast adjustments were made by ImageJ (National Institutes of Health).

### Microglia ablation

Microglial ablation was achieved through the administration of a chow containing the colony-stimulating factor 1 receptor (CSF1R) inhibitor, PLX3397 (600 mg/kg, Chemgood), provided *ad libitum* for a minimum of 2 weeks to ensure complete depletion of microglia. The efficiency of microglial ablation was evaluated through immunostaining of brain sections.

### Microglia dynamics and morphology analyses

Two-photon z-stacked and time-lapse images were aligned using the ImageJ plugin, StackReg, to correct any image shifts. For the analysis of microglia dynamics in two-photon images, the cumulative area occupied by microglia was quantified over time. Firstly, a z-projected image was thresholded using the Li method in ImageJ. Cumulative maximum projections of the microglial signal from time (t_x_) and previous time points (t_0_ + t_1_ + … + t_x_) were generated, and the area filled by the microglial signal was calculated from the beginning (t0) to the end of the imaging session (t_0_ + t_1_ + … + t_30_). Subsequently, the data were normalized to the area filled at “t_0_” to obtain a relative area surveyed over time, independent of the initial microglia morphology.

For the analysis of microglia morphology from two-photon images, a z-projected image was thresholded using the Li method in ImageJ. All microglia with clear soma were manually selected for Sholl analysis. Additionally, to analyze microglia morphology from Iba1 immunostaining, a z-projected image was initially segmented using the trainable Weka Segmentation plug-in in ImageJ. Subsequently, 10-15 microglia per mouse were randomly selected for Sholl analysis.

### Microglia-neuron soma interaction analyses

Two-photon Z-stacked time-lapse images were aligned using the ImageJ plugin, HyperStackReg, to correct any image shifts. To quantify *in vivo* interactions between neuronal somata and microglial processes, three planes of two-photon images (4 μm thick) were average projected. A segmented line (1.5 μm wide) was then drawn along the neuron soma. Subsequently, the “Plot Profile” tool was employed to profile the GFP fluorescence intensity (representing microglia processes) along the tdTomato^+^ neuron soma. All neurons with clear soma were included in the analysis, excluding those interacting with microglia soma. The raw fluorescence intensity was converted into ΔF/F_0_, where F_0_ was defined as the 50^th^ percentile value for each neuron. Next, all ΔF/F_0_ values were normalized by the maximum ΔF/F_0_ observed across all analyzed neurons in this time-lapse average projected image. Instances with a relative ΔF/F_0_>0.3 for more than 0.5 μm were considered as interaction events. The number of interaction events during the 30-minute imaging period and the contact duration were then calculated.

For the analysis of microglia-neuron soma interaction in immunostaining images, a segmented line (1.5 μm wide) was drawn along the Kv2.1^+^ neuron soma. A single plane of confocal images was split into a Kv2.1 channel and a P2Y12 channel. Similar to the two-photon image analysis, the “Plot Profile” tool was used to profile identical segmented lines in each channel. F_0_ was defined as the 5^th^ percentile value for Kv2.1 and the 50^th^ percentile value for P2Y12. Data processing steps were the same as the analysis of microglia-neuron soma interaction in two-photon images. 50 neurons per mouse were randomly selected for this analysis.

### Neuronal calcium imaging analyses

Two-photon time-lapse images underwent registration using the ImageJ plugin, TurboReg, to correct any image shifts. For the analysis of neuronal calcium activity, an average intensity image of the entire video was generated to facilitate the selection of neuron somata. Neuron somata were manually drawn using the oval selection tool. Neurons lacking “donut-shaped” calcium activity were excluded from the analysis. Then, mean fluorescent intensity values were obtained and subsequently converted into ΔF/F_0_. The baseline fluorescence was defined as the lower 25^th^ percentile value across all frames. Neurons exhibiting ΔF/F_0_>0.25 were considered “active”. Parameters such as neuronal active time, signal area, and calcium transient events and duration were calculated accordingly.

To evaluate the neural network synchronization, time-lapse calcium imaging data were analyzed using the MATLAB-based Mic2net program.^27^ Mic2net algorithms provide automated processing and quantitative assessment of neuronal calcium imaging data that has been pre-processed using ImageJ. Cross-correlation analysis was employed to determine the synchronization between pairs of neurons. To rule out insignificant correlations, cut-off values were determined using a scrambled data set. This set was generated by shuffling the time series of calcium imaging to random starting points. The 99th percentile of the cross-covariance values from the scrambled data set was then applied as the cut-off value. Connectivity was defined as the number of neurons exhibiting a correlation coefficient higher than the cut-off, divided by the total number of neurons. Connectance, on the other hand, was defined as the number of significant edges divided by the maximum possible number of edges in the network, providing a quantitative measure of the overall network synchronization.

### Single-cell RNA sequencing database queries

The Tabula Muris database served as the resource for determining *Tmem119 and Csf1r* expression across 20 mouse organs at the single-cell level.^58^ All indexed R objects (https://figshare.com/articles/dataset/Robject_files_for_tissues_processed_by_Seurat/5 582126) were updated to the newest *Seurat v4.0* object^59^ using the UpdateSeuratObject function. These objects were then combined into one *Seurat* object using the *merge* function. Normalization of gene counts was performed by *NormalizeData* function. The data were further scaled by *ScaleData* function. Highly variable features were called by *FindVariableFeatures* function. Initial dimensionality reduction was carried out via principal component analysis (PCA), followed by t-distributed stochastic neighbor embedding (tSNE). Visualization of the results was achieved using *Dimplot* for tSNE plots. *FeaturePlot* function was employed to illustrate *Tmem119 and Csf1r* expression levels.

### Statistics

Detailed statistical information, including sample size and statistical methods, is provided in the figure legends corresponding to each specific experiment. Generally, normality was first assessed, and all data followed the Gaussian distribution. An unpaired 2-tailed *t-*test was employed for comparing experiments involving 2 groups. In instances where 2 time points from the same animal were compared, a paired 2-tailed *t-*test was used. For comparisons with more than 2 time points from the same animal, repeated measures one-way ANOVA was performed, followed by a Bonferroni’s *post hoc* analysis for multiple comparisons. Results are presented as the mean ± standard error of the mean (SEM), and statistical significance was determined when *p*<0.05. Statistical analyses were performed using GraphPad Prism 10 software. Experimental designs and sample sizes were determined to minimize animal usage and distress while ensuring sufficiency for detecting robust effect sizes.

## Data availability

All data reported in this paper will be shared by the lead contact upon request.

## Code availability

All original codes are deposited on GitHub and will be publicly available as of the date of publication.

## Author contributions

S.Z. and L.-J.W. designed experiments and wrote the manuscript. S.Z. performed most of the experiments. L.W. processed and performed de novo analyses of existing RNA sequencing datasets. Y.L. and J.Z. performed some experiments. S.Z. analyzed data. A.D.U. and L.-J.W. proofread the manuscript.

## Acknowledgments

We thank members of the Wu lab for insightful discussions. This work was supported by the following grants from the National Institutes of Health: R35NS132326 (L.-J.W.) and K99NS126417 (A.D.U.).

## Supplemental Videos

**Video S1. Chemogenetic activation of microglial Gi signaling results in microglial process retraction in awake mice (related to Figure 1).** *In vivo* two-photon time-lapse imaging of microglial dynamics before and 5-60 minutes after DCZ injection in *Tmem119^CreER/+^:R26^hM4Di/+^:Cx3cr1^GFP/+^*mice. GFP^+^ microglia (green) retract processes after DCZ injection. Scale bar: 10 μm.

**Video S2. Activation of microglial Gi-DREADD leads to microglial process retraction in anesthetized mice (related to Figure 1).** *In vivo* two-photon time-lapse imaging of microglial dynamics before and 5-60 minutes after DCZ injection in anesthetized *Tmem119^CreER/+^:R26^hM4Di/+^:Cx3cr1^GFP/+^* mice. GFP^+^ microglia (green) retract processes after DCZ injection. Scale bar: 10 μm.

**Video S3. Activation of microglial Gi signaling leads to fewer interactions between microglia processes and neuron somata (related to Figure 3).** *In vivo* two-photon time-lapse imaging of microglia-neuron soma interactions before and 0-60 minutes after DCZ injection in *Tmem119^CreER/+^:R26^hM4Di/+^:Cx3cr1^GFP/+^*mice. Arrows indicate the retracting microglial processes (GFP^+^, green) from neuronal soma (tdTomato^+^, red). Scale bar: 5 μm.

## Supplemental figures

**Figure S1.**
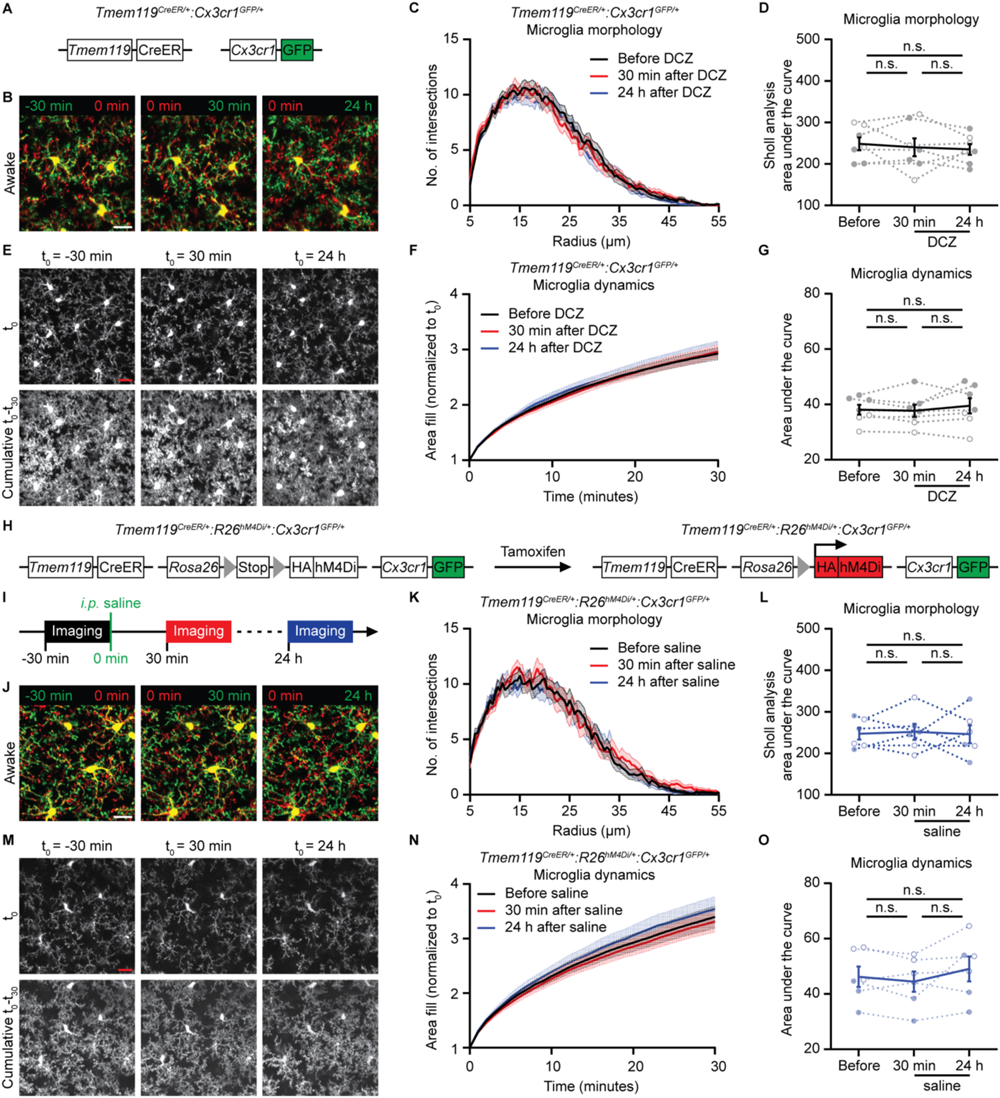
Saline injection does not affect microglia surveillance in Tmem119^CreER/+^:R26^hM4Di/+^:Cx3cr1^GFP/+^ mice (related to Figure 1). **(A)** The genetic strategy of *Tmem119^CreER/+^:Cx3cr1^GFP/+^* control mice lacking Gi-DREADD expression. **(B)** Representative two-photon images of microglia in control mice at each study phase. Left: before (green: -30 min; red: 0 min). Mid: 30 minutes after (red: 0 min; green: 30 min). Right: 24 h after (red: 0 min; green: 24 h). Scale bar: 20 μm. (**C** and **D**) Sholl analysis (C) and the area under the curve quantification (D) reveal no morphology alteration in microglia after DCZ administration in control mice (33 microglia from N=7 mice in each group). **(E)** Two-photon images of cumulative microglial surveillance before, 30 minutes after, and 24 hours after DCZ administration in control mice. Upper: microglial surveillance area at the beginning of each study phase. Lower: 30-minute cumulative microglial surveillance area of each study phase. Scale bar: 20 μm. (**F** and **G**) Normalized cumulative microglial surveillance area (F) and the area under the curve quantification (G) show unchanged microglial surveillance after DCZ administration in control mice (N=7 mice in each group). **(H)** The genetic strategy employed in *Tmem119^CreER/+^:R26^hM4Di/+^:Cx3cr1^GFP/+^* mice for the expression of hM4Di (Gi-DREADD) and GFP in microglia. **(I)** Timeline of *in vivo* two-photon imaging in awake mice. 30-minute videos were recorded before, 30 minutes, and 24 hours after saline injection. **(J)** Representative two-photon images of microglia morphology at each study phase in *Tmem119^CreER/+^:R26^hM4Di/+^:Cx3cr1^GFP/+^* mice. Left: before (green: -30 min; red: 0 min). Mid: 30 minutes after (red: 0 min; green: 30 min). Right: 24 h after (red: 0 min; green: 24 h). Scale bar: 20 μm. (**K** and **L**) Sholl analysis (K) and the area under the curve quantification (L) demonstrate no significant microglial morphology alteration after saline administration in *Tmem119^CreER/+^:R26^hM4Di/+^:Cx3cr1^GFP/+^* mice (24 microglia from N=6 mice in each group). (**M**) Two-photon images of cumulative microglial surveillance before, 30 minutes after, and 24 hours after saline administration in *Tmem119^CreER/+^:R26^hM4Di/+^:Cx3cr1^GFP/+^*mice. Upper: microglial surveillance area at the beginning of each study phase. Lower: 30-minute cumulative microglial surveillance area of each study phase. Scale bar: 20 μm. (**N** and **O**) Normalized cumulative microglial surveillance area (N) and the area under the curve quantification (O) illustrate unchanged microglia surveillance after saline administration in *Tmem119^CreER/+^:R26^hM4Di/+^:Cx3cr1^GFP/+^*mice (N=6 mice in each group). In all graphs, each point indicates an individual mouse (solid dots: male mice; hollow dots: female mice). Data are represented as the mean ± SEM. Repeated measures one-way ANOVA followed by a Bonferroni’s *post hoc* test (D, G, L, and O). n.s.: not significant.

**Figure S2.**
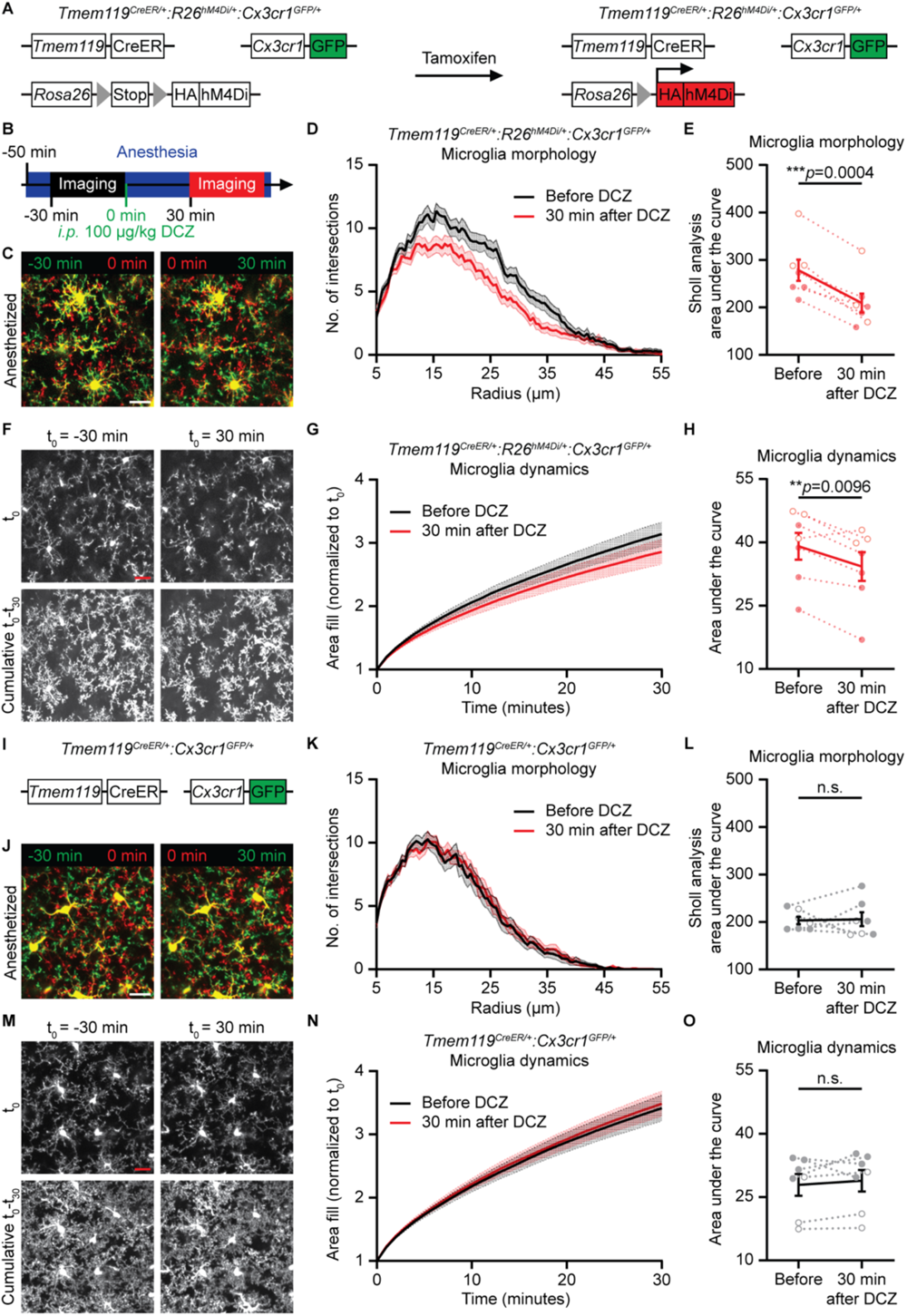
Microglia decreased process surveillance after chemogenetic activation of microglial Gi signaling in anesthetized mice (related to Figure 1). (**A**) The genetic strategy employed in *Tmem119^CreER/+^:R26^hM4Di/+^:Cx3cr1^GFP/+^* mice for the expression of hM4Di (Gi-DREADD) and GFP in microglia. (**B**) Timeline of *in vivo* two-photon imaging in anesthetized mice. 20 minutes after general anesthesia induction, a 30-minute baseline video was recorded. Immediately after baseline imaging, 100 μg/kg DCZ was *i.p.* administered, and a 30-minute video was acquired 30-60 minutes post-injection. (**C**) Representative two-photon images of microglia morphology at each study phase in anesthetized *Tmem119^CreER/+^:R26^hM4Di/+^:Cx3cr1^GFP/+^*mice. Increased red signal area in the left panel suggests microglial process extension during anesthesia, while increased red signal area in the right panel indicates microglial process retraction after DCZ injection. Left: before (green: -30 min; red: 0 min). Right: 30 minutes after (red: 0 min; green: 30 min). Scale bar: 20 μm. (**D** and **E**) Sholl analysis (D) and the area under the curve quantification (E) reveal microglia process retraction 30 minutes after DCZ administration in anesthetized *Tmem119^CreER/+^:R26^hM4Di/+^:Cx3cr1^GFP/+^* mice (33 microglia from N=7 mice in each group). **(F)** Two-photon images of cumulative microglial surveillance before and 30 minutes after DCZ administration in anesthetized *Tmem119^CreER/+^:R26^hM4Di/+^:Cx3cr1^GFP/+^*mice. Microglia exhibit reduced surveillance area following DCZ injection. Upper: microglial surveillance area at the beginning of each study phase. Lower: 30-minute cumulative microglial surveillance area of each study phase. Scale bar: 20 μm. (**G** and **H**) Normalized cumulative microglial surveillance area (G) and the area under the curve quantification (H) illustrate decreased microglial surveillance 30 minutes after DCZ administration in anesthetized *Tmem119^CreER/+^:R26^hM4Di/+^:Cx3cr1^GFP/+^* mice (N=7 mice in each group). **(I)** The genetic strategy of *Tmem119^CreER/+^:Cx3cr1^GFP/+^* control mice lacking Gi-DREADD expression. **(J)** Representative two-photon images of microglia morphology at each study phase in anesthetized control mice. Left: before (green: -30 min; red: 0 min). Right: 30 minutes after (red: 0 min; green: 30 min). Scale bar: 20 μm. (**K** and **L**) Sholl analysis (K) and the area under the curve quantification (L) reveal no significant microglia morphology alteration after DCZ administration in anesthetized control mice (32 microglia from N=7 mice in each group). (**M**) Two-photon images of cumulative microglial surveillance before and 30 minutes after DCZ administration in anesthetized control mice. Upper: microglial surveillance area at the beginning of each study phase. Lower: 30-minute cumulative microglial surveillance area of each study phase. Scale bar: 20 μm. (**N** and **O**) Normalized cumulative microglial surveillance area (N) and the area under the curve quantification (O) demonstrate unchanged microglial surveillance after DCZ administration in anesthetized control mice (N=7 mice in each group). In all graphs, each point indicates an individual mouse (solid dots: male mice; hollow dots: female mice). Data are represented as the mean ± SEM. Paired *t*-test (E, H, L, and O). n.s.: not significant. \*\**p*<0.01 and \*\*\**p*<0.001. The exact *p* values are directly provided in the figure panels.

**Figure S3.**
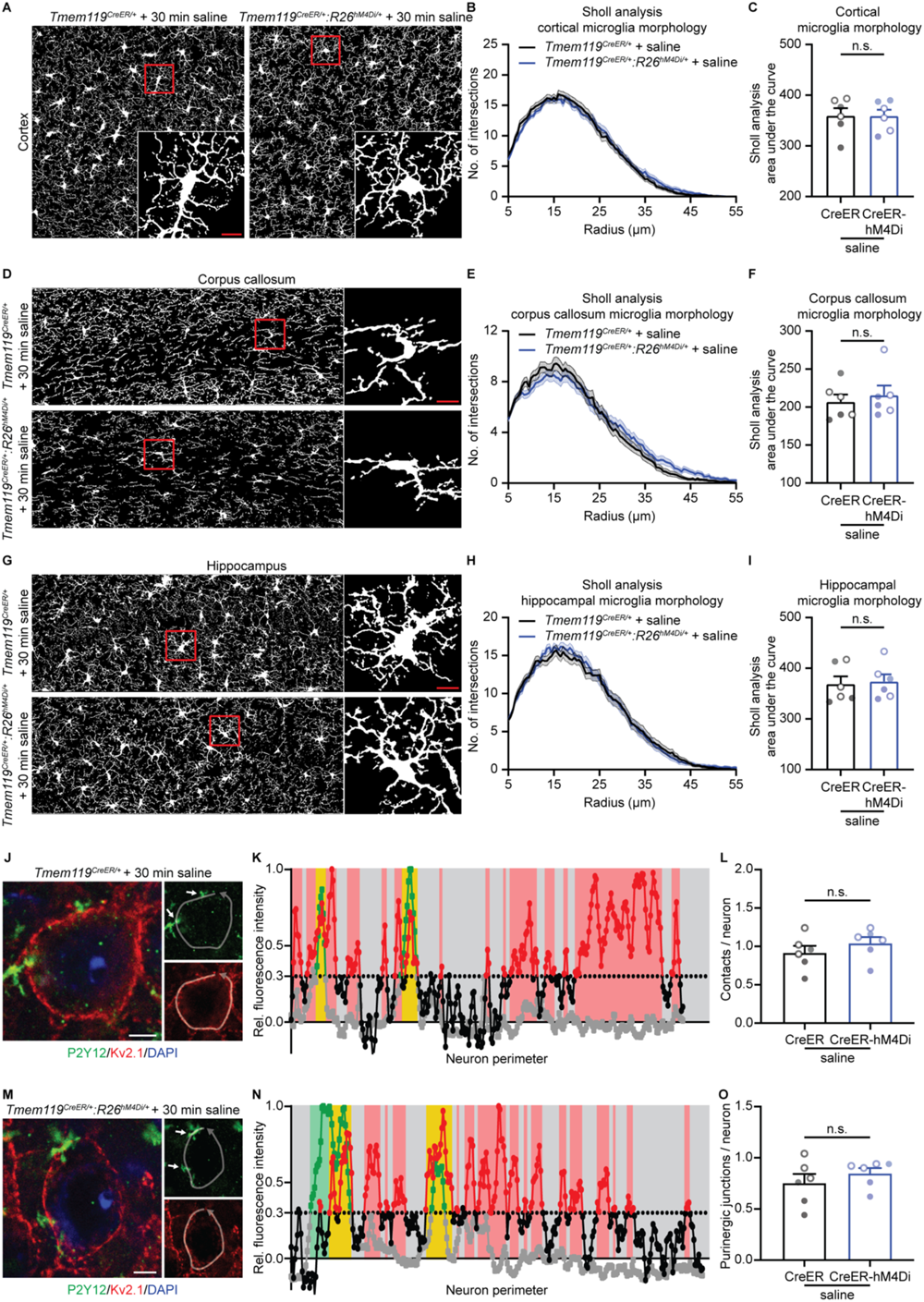
Saline injection does not affect microglia process length in various brain regions and microglia-neuron soma interactions in *Tmem119^CreER/+^:R26^hM4Di/+^* mice (related to Figure 2 and Figure 3). (**A**-**C**) Analysis of cortical microglia morphology in saline-injected *Tmem119^CreER/+^* and *Tmem119^CreER/+^:R26^hM4Di/+^* mice. Representative Iba1 mask (A), Sholl analysis (B), and the area under the curve quantification (C) reveal no significant changes in cortical microglia processes (60 microglia from N=6 mice in each group). (**D**-**F**) Evaluation of microglial morphology in the corpus callosum. Representative Iba1 mask (D), Sholl analysis (E), and the area under the curve quantification (F) demonstrate that saline injection did not induce morphological alterations in the corpus callosum microglia of *Tmem119^CreER/+^* and *Tmem119^CreER/+^:R26^hM4Di/+^* mice (60 microglia from N=6 mice in each group). (**G**-**I**) Assessment of hippocampal microglia. Representative Iba1 mask (G), Sholl analysis (H), and the area under the curve quantification (I) indicate that hippocampal microglia morphology remained unchanged after saline injection in *Tmem119^CreER/+^*and *Tmem119^CreER/+^:R26^hM4Di/+^* mice (60 microglia from N=6 mice in each group). (**J**-**N**) Immunostaining (J and M) and plots of neuron perimeter versus fluorescence intensity (K and N) showing neuronal Kv2.1 membrane domains (Kv2.1^+^ and P2Y12^neg^, red), non-purinergic microglial contacts (Kv2.1^neg^ and P2Y12^+^, green), and purinergic junctions (Kv2.1^+^ and P2Y12^+^, yellow). 30 minutes after saline injection, neurons had comparable microglial contacts and purinergic junctions in both groups. Arrows indicate the purinergic junction (J and M). (**L** and **O**) Quantifications revealing microglia processes had similar total contacts (L) and purinergic junctions (O) per neuron 30 minutes after saline administration in both groups (300 neurons from N=6 mice in each group). In all graphs, each point indicates an individual mouse (solid dots: male mice; hollow dots: female mice). Data are represented as the mean ± SEM. Unpaired *t*-test (C, F, I, L, and O). n.s.: not significant. Scale bar: 10 μm (A, D, and G); 5 μm (J and M).

**Figure S4.**
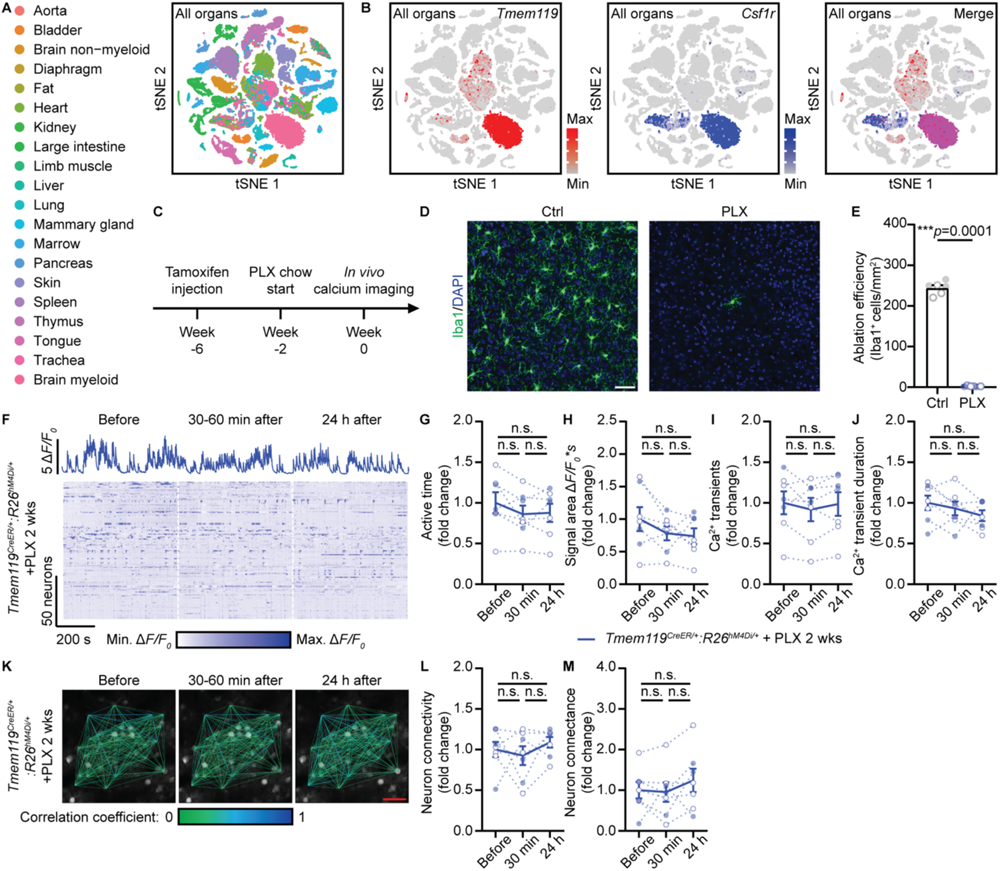
Microglia ablation rescues microglial Gi signaling-induced neuronal hypoactivity and hyposynchronization (related to Figure 4). **(A)** Query of Tabula Muris database showing cells in 20 mouse organs by t-SNE plot. **(B)** t-SNE plots of all cells from the database demonstrating the dense expression of *Tmem119* and *Csf1r* in brain myeloid cells and sparse detection in other peripheral organs. **(C)** Timeline depicting the induction of Gi-DREADD expression, microglia ablation, and *in vivo* two-photon calcium imaging for *Tmem119^CreER/+^* and *Tmem119^CreER/+^:R26^hM4Di/+^*mice. (**D** and **E**) Representative images (D) and quantitative analysis (E) demonstrating efficient microglia ablation (Iba1^+^, green) in the brain with PLX treatment (N=6 mice in each group). Scale bar: 50 μm. (**F**) Single neuronal calcium activity traces (upper) and heatmap of all neuronal calcium activity (lower) before, 30-60 minutes after, and 24 hours after DCZ administration in microglia-ablated *Tmem119^CreER/+^:R26^hM4Di/+^*mice. No neuronal activity alteration was observed after DCZ injection. (**G**-**J**) Quantifications revealing neuronal active time (G), signal area (H), calcium transient events (I), and calcium transient duration (J) after the DCZ injection in *Tmem119^CreER/+^:R26^hM4Di/+^* mice (139 neurons from N=7 mice in each group). **(K)** Representative neuronal network analysis plotted on top of two-photon calcium images in microglia ablated *Tmem119^CreER/+^:R26^hM4Di/+^*mice at each study phase. No alteration was observed after DCZ injection. Scale bar: 50 μm. (**L** and **M**) Quantifications revealing unaltered neuron connectivity (L) and neuron connectance (M) after the DCZ injection in microglia ablated *Tmem119^CreER/+^:R26^hM4Di/+^*mice (N=7 mice in each group). In all graphs, each point indicates an individual mouse (solid dots: male mice; hollow dots: female mice). Data are represented as the mean ± SEM. Unpaired *t*-test (E). Repeated measures one-way ANOVA followed by a Bonferroni’s *post hoc* test (G-J, L, and M). n.s.: not significant. \*\*\**p*<0.001. The exact *p* values are directly provided in the figure panels.

